# Robust perisomatic GABAergic self-innervation inhibits basket cells in the human and mouse supragranular neocortex

**DOI:** 10.1101/760983

**Authors:** Viktor Szegedi, Melinda Paizs, Judith Baka, Pal Barzo, Gabor Molnar, Gabor Tamas, Karri Lamsa

## Abstract

Inhibitory autapses are self-innervating synaptic connections in GABAergic interneurons in the brain. Autapses in neocortical layers have not been systematically investigated, and their function in different mammalian species and specific interneuron types is poorly known. We investigated GABAergic parvalbumin-expressing basket cells (pvBCs) in layer 2/3 (L2/3) in mice as well as in human neocortical tissue resected in deep-brain surgery. Most pvBCs showed robust GABA_A_R-mediated self-innervation in both species, but autapses were rare in nonfast spiking GABAergic interneurons. Light- and electron microscopy analyses revealed pvBC axons innervating their own soma and proximal dendrites. GABAergic self-inhibition conductance was similar in human and mouse pvBCs and comparable to that of synapses from pvBCs to other L2/3 neurons. Autaptic conductance prolonged somatic inhibition in pvBCs after a spike and inhibited repetitive firing. Perisomatic autaptic inhibition has evolved in pvBCs of various cortical layers and different mammalian species to control discharge of these interneurons.

## INTRODUCTION

Autapses are synapses made by a neuron onto itself (Bekkers JM 2003; Deleuze C et al. 2014). Although studies on experimental animals have reported autaptic self-innervation in some inhibitory as well as in excitatory neurons in the brain, (Karabelas AB and DP Purpura 1980; Lubke J et al. 1996; Thomson AM et al. 1996; Cobb SR et al. 1997; Pouzat C and A Marty 1998; Bacci A et al. 2003; Connelly WM and G Lees 2010; Karayannis T et al. 2010; Jiang M et al. 2015; Yin L et al. 2018), little is known about autapses in human nerve cells, and only a single study has reportedly investigated the operation of autaptic self-inhibition in the human neocortex (Jiang M et al. 2012). Hence, how autaptic inhibitory circuits operate in the human brain compared to those in common experimental animals remains largely unknown.

Few notable studies in rodents have demonstrated GABAergic autapses in neocortical deep layer fast-spiking parvalbumin-expressing basket cells (pvBCs) by showing functional and pharmacological evidence for self-inhibition. In layer 5 of the infragranular neocortex, GABA_A_R-mediated self-inhibition prolongs the pvBC spiking interval during sustained high-frequency firing (Bacci A and JR Huguenard 2006; Manseau F et al. 2010). Evidence for autapses also exists in the human epileptic infragranular neocortex, where high-frequency spike bursts are associated with autaptic GABA release from fast-spiking interneurons (Jiang M et al. 2012). However, the operation of autaptic self-inhibition in superficial neocortical layers is unknown (Tamas G et al. 1997).

GABAergic inhibition at the perisomatic region efficiently controls spike output (Tremblay R et al. 2016; Feldmeyer D et al. 2018). Hence, pvBCs are key players synchronizing neuronal network activity through their inhibitory synapses (Cobb SR et al. 1995; Pouille F and M Scanziani 2001; Wood KC et al. 2017; Cardin JA 2018), and altered pvBC firing is often linked to pathological network activities (Jiang X et al. 2016) (Palop JJ and L Mucke 2016; Dienel SJ and DA Lewis 2018). In addition, perisomatic inhibition through autapses can efficiently regulate pvBC firing (Bacci A and JR Huguenard 2006; Guo D et al. 2016; Yilmaz E et al. 2016). Computational models can help us understand the role of pvBC self-inhibition in the generation and maintenance of cortical network activities (Connelly WM 2014; Guo D et al. 2016; Yilmaz E et al. 2016), but very little is known about autapses themselves, including their occurrence and inhibitory efficacy in distinct interneuron types.

We investigated GABAergic autapses in human and mouse supragranular layer 2/3 pvBCs and some nonfast spiking interneurons. We show that GABA_A_R-mediated inhibition is present in most pvBCs in both species and that perisomatic autaptic contacts in these interneurons suppress excitability and inhibit repetitive firing. Autapses are rare in nonfast spiking GABAergic interneurons. We conclude that GABAergic interneuron autapses are a standard and cell type-specific microcircuit feature in the mammalian neocortex that mediates strong perisomatic self-inhibition of pvBCs.

## RESULTS

We investigated autapses in 45 supragranular layer 2/3 (L2/3) pvBCs and 22 nonfast spiking interneurons (nonFSINs) in human neocortical tissue resected from frontal, temporal or occipital areas during deep brain surgery to obtain access to subcortical pathological targets (tumor, cyst, aneurysm or catheter implant). For comparison, we studied pvBC in the mouse somatosensory cortex. Supplementary Table 1 lists human interneurons and shows their spike kinetics, immunohistochemical reactions and details of the cortical human tissue material.

### Autaptic GABA_A_R-mediated self-innervation in human supragranular pvBCs

First, we recorded cells in whole-cell current clamp using elevated intracellular chloride (75 mM) that makes a GABA_A_R-mediated potential robustly depolarizing (Fig. 1a_i_) (Bacci A et al. 2003; Bacci A and JR Huguenard 2006). Under these conditions, unitary spikes (interval 10 s) in 11 of the 13 pvBCs studied triggered GABA_A_R-mediated depolarizing potentials (5.44 ± 0.73 mV peak amplitude, n = 11, SWP = 0.380), showing a 0.92 ± 0.04 ms onset delay to the action potential peak (n = 11, SWP = 0.132, see Fig. 1a_ii_). In contrast, nonFSINs showed GBZ-sensitive autaptic potentials in only 1 of the 14 cells studied (action inward current width 0.886 ± 0.036 ms vs. 0.536 ± 0.024 ms in pvBCs, P < 0.001, t-test).

**Figure 1.**
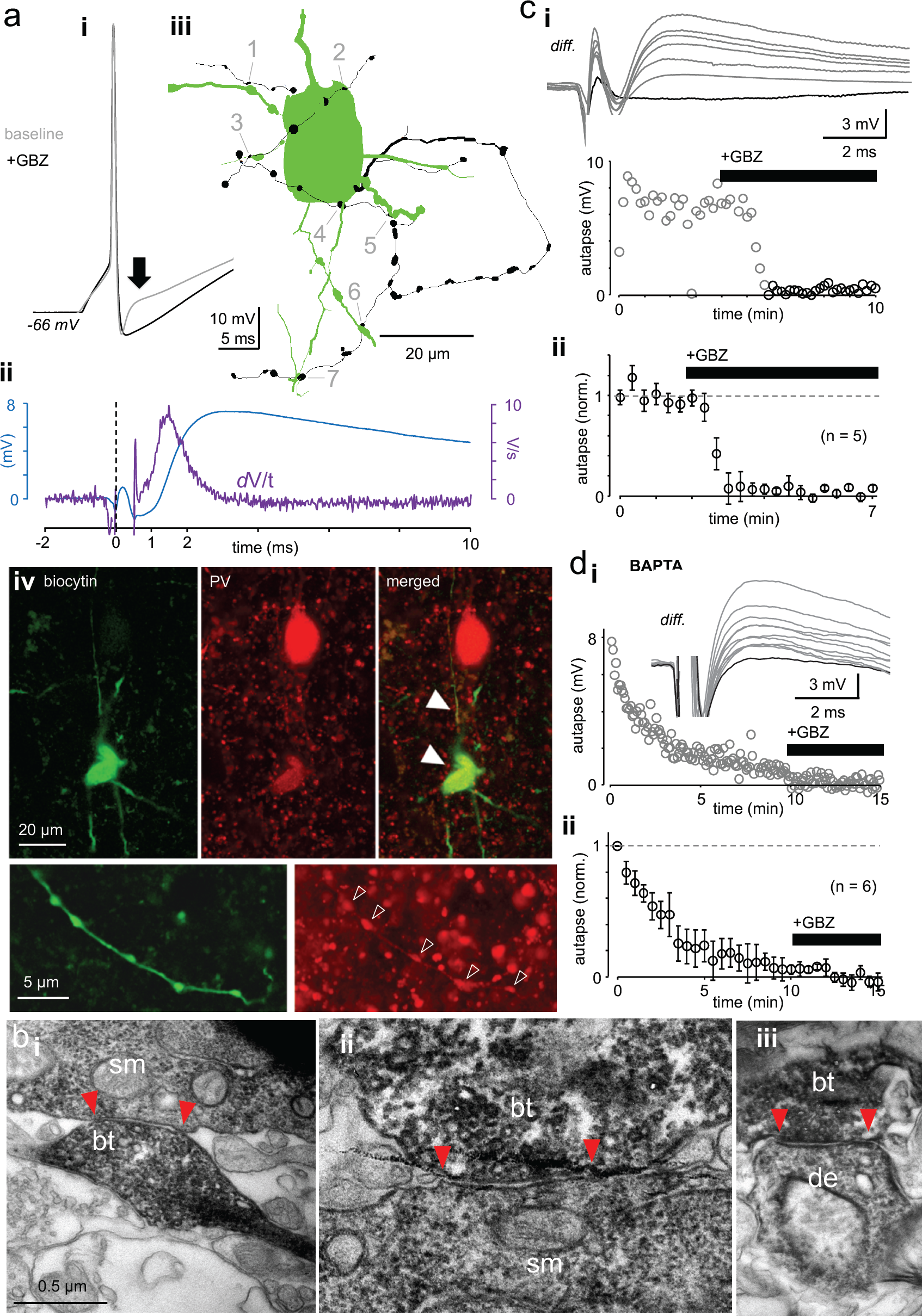
Autaptic GABA_A_R-mediated innervation in human supragranular pvBCs. **a)** Elevated intracellular chloride reveals GABA_A_R-mediated self-innervation in layer 2/3 parvalbumin-immunopositive basket cells (pvBCs). (i) Gray trace shows unitary spike-evoked depolarizing autaptic potential (peak indicated by arrow) measured in current clamp. Black trace shows its blockage by gabazine (GBZ, 10 μM). Traces are averages of six. (ii) Autaptic potential (blue trace) shown as a differential of the response at baseline and in the presence of GBZ. The first derivate (dV/t) of this potential reveals < 1 ms onset delay to the action potential peak (at 0 time point). (iii) Light microscopic reconstruction of the same cell reveals a self-innervating axon (black), forming boutons (1-7) in close apposition to its own soma and proximal dendrites. (iv) Confocal fluorescence images of the same cell (biocytin-Alexa488) show pv (Cy3) immunopositive soma, dendrites (solid arrowheads) and axon (open arrowheads). **b)** Innervation pattern of autaptic boutons. Electron microscopic images illustrate a sample biocytin-filled pvBC axon forming autaptic appositions to the soma and proximal dendrite. (i-ii) Sample autaptic boutons (*bt*) terminating on soma (*sm*). (iii) A bouton forming autaptic contact in the same cell to proximal dendrite (*de*). Red arrows indicate the apposition zone. **c)** Autaptic response is systematically blocked by gabazine (GBZ, 10 μM). (i) *Top:* Unitary spike-evoked autaptic potentials shown as differential as in aii (*diff.*). Traces are during baseline and show the effect of GBZ wash-in (interval 10 s). Black trace shows full blockade. *Bottom:* Plot shows the GBZ effect on the autaptic potential peak amplitude in the same experiment. (ii) Mean ± se of 5 similar experiments (bin 30 s, amplitude baseline-normalized). **d)** Intracellular BAPTA (10 mM) abolishes autaptic potential gradually. (i) Slow inhibition of a spike-evoked autaptic potential peak amplitude in one experiment (interval 5 s). GBZ (10 μM) was applied at the end. *Inset*: Traces show vanishing autaptic potential (*diff.*) by course of experiment with BAPTA-containing filling solution. The black trace is in the presence of GBZ. (ii) Mean ± se of 6 similar experiments (30 s bin, amplitude normalized by the average of the first 30 s).

We visualized recorded cells (filled with biocytin) with fluorophore-streptavidin (Fig. 1a_iii_) and studied their parvalbumin immunoreactivity using confocal fluorescence microscopy (Fig. 1a_iv_) (Supplementary Table 1). In some cells with fully recovered intact soma, we investigated anatomical evidence for self-innervation after avidin-biotinylated horseradish peroxidase reaction. By using a light microscopic study of somatic area (in 50 μm-thick section) (n = 5 cells), we found apparent self-innervation showing close apposition boutons (range 3-8 per cell) formed by their labeled axons to the proximal dendrites (distance to soma = 30 μm, 11 – 46 μm) (median, quartiles, n = 17 in 5 cells) and on the boundaries of soma (range 2-5 per soma, n = 10 in 3 cells). Because dense peroxidase products mostly obscured the observation of close appositions on the soma (Tamas G et al. 1997), we specifically looked for somatic autaptic junctions using electron microscopy in separate pvBCs. The analysis in two cells showing electrophysiological evidence for GABAergic autapses revealed pvBC biocytin-filled axon terminal boutons forming contacts to their own soma (1 and 4 contacts) and proximal dendrites (1 and 2 contacts), revealing postautaptic density in the neuron (Fig. 1b_i-iii_).

The autaptic response was readily blocked by GBZ (10 μM) (P <0.001, n = 5) (Fig. 1c_i,ii_) or gradually abolished by intracellular calcium-chelator BAPTA (20 mM), which suppresses action potential-dependent vesicle release (n = 6, P <0.001) (Wilcoxon signed rank test) (Fig. 1d_i,ii_) (Bacci A et al. 2003). The autaptic response amplitude and onset delay values were measured by subtracting spike-elicited responses at baseline and in the presence of GABA_A_R blocker gabazine (GBZ, 10 μM) (see Fig. 1a_ii_).

### GABAergic self-inhibition strength in human and mouse pvBCs evoked by unitary spikes

Next, we studied autaptic self-innervation conductance (G_aut_) and its inhibitory strength in pvBCs. We used voltage clamp with close-to-physiological intracellular (8 mM) chloride (Verheugen JA et al. 1999; Connelly WM and G Lees 2010) to minimize error in GABA_A_R- mediated conductance emerging from artificial transmembrane chloride gradient (Hille B 2001). Spike-evoked autaptic outward current (interval 10 s, recorded at −43 to −55 mV) was uncovered by subtracting response in gabazine (average of 6 after wash-in of GBZ 10 μM) from baseline responses (Fig. 2a_i-iii_). In human pvBCs, a spike-evoked autaptic response showed peak G_aut_ of 3.79 nS, 2.39-6.45 nS (median, quartiles, n = 14 cells) and 4.72 ± 0.27 ms decay time constant (SW P = 0.782, n = 10 cells, defined in cells showing the largest autaptic current) (monoexponential curve fitting r2 range from 0.86 to 0.97). The autaptic current peak amplitude had a 2.57± 0.14 ms (SW P = 0.55) delay to the action inward current peak (n = 14).

**Figure 2.**
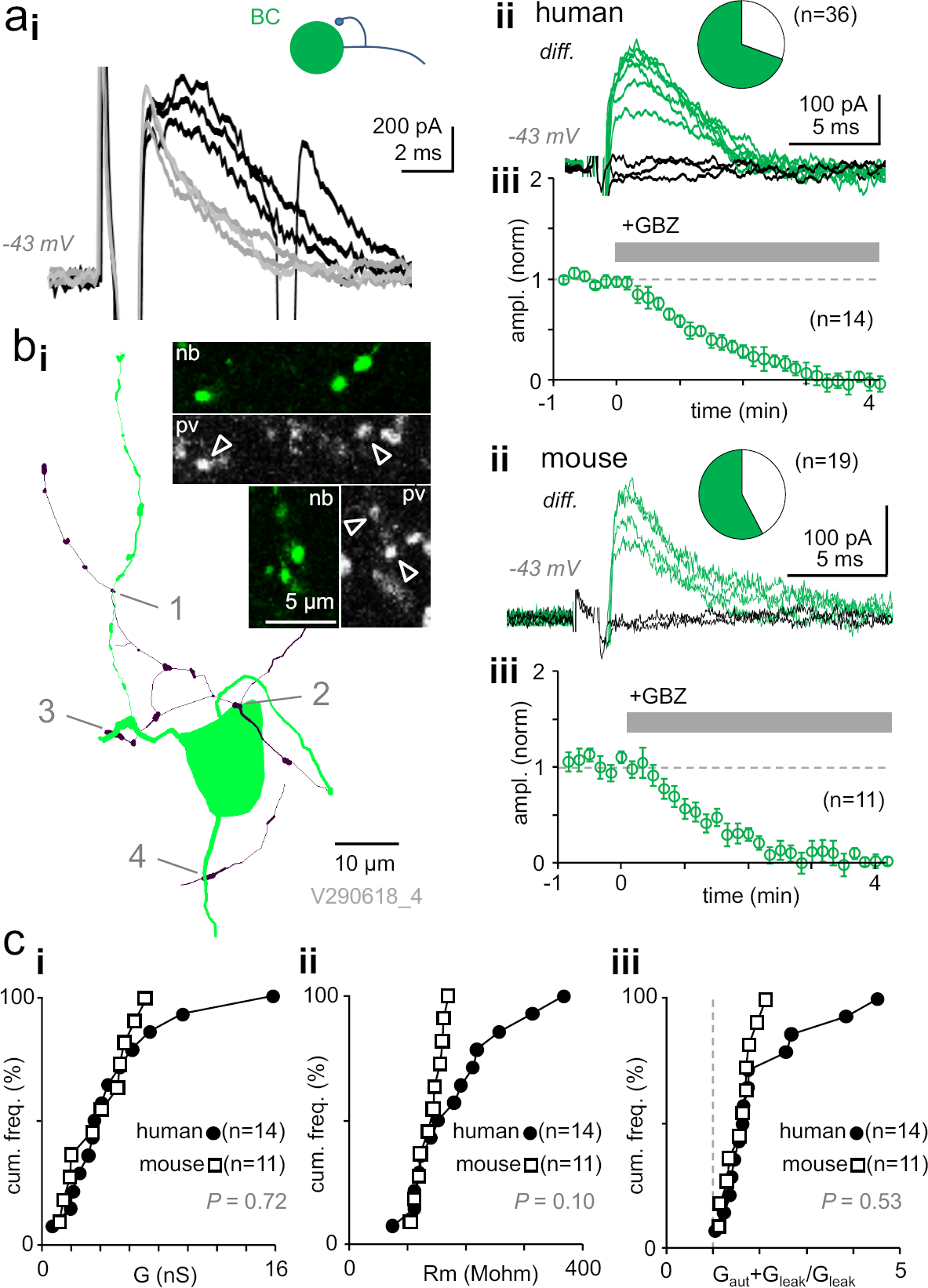
Autaptic self-inhibition strength in human and mouse pvBCs evoked by unitary spikes. **a)** A voltage clamp recording in human pvBCs shows autaptic GABA_A_R-mediated inhibitory current (intracellular chloride 8 mM). (i) Unitary spike-evoked responses in a human pvBC during baseline (black) and after wash-in of gabazine (GBZ, 5 min, gray) (interval 10 s, at −43 mV). Note the second action potential escape current generated occasionally. (ii) Subtraction of the traces reveals GABAergic current. *Top:* Traces show the autaptic GBZ-sensitive current (*diff.*) in baseline (green) and when fully blocked (black, subtraction of traces in GBZ). Inset pie chart shows the proportion of cells with GBZ-sensitive autapse (green) in all pvBCs studied (in 25 of 36). (iii) Mean ± sem of GBZ wash-in in all pvBCs (n = 14, amplitude baseline-normalized, 30 s bin). **b)** GABA_A_R-mediated self-innervation and autaptic current in mouse pvBCs. (i) A visualized mouse pvBC perisomatic area showing axon (black) boutons forming close appositions (1-4) to its own soma and proximal dendrites. Confocal microscopic micrographs illustrate the pv immunopositive (pv, Cy5) axon of the cell (nb, Alexa488). (ii) GBZ-sensitive autaptic current (*diff*) at baseline (green). Black traces are the subtraction of traces in GBZ. Inset pie chart shows the proportion of mouse pvBCs with autapses (in 11 of 19 cells). (iii) Mean ± sem of 11 experiments showing GABAergic current blockade by wash-in of GBZ (amplitude baseline-normalized). **c)** Autaptic self-inhibition efficacy in human and mouse pvBCs. Cumulative presentation of solid and open symbols indicates the average in cells. (i) Self-inhibition peak conductance (G) in human and mouse pvBCs. (ii) Input resistance (R_m_) in humans and mice. (iii) Calculated total input conductance of a cell during the autaptic current peak (G_aut_ + G_leak_), divided by its resting conductance (G_leak_). The ratio shows how much self-inhibition increases membrane leakage.

Altogether, experiments including the current- and voltage clamp measurements above confirmed GBZ-sensitive autapses in 25 of 36 (69.4 %) human pvBCs studied (see Fig. 2a_ii_).

On average, the human pvBC autapse peak conductance was not different from synaptic conductance (G_syn_) tested in 20 monosynaptically connected pvBC-PC pairs (Supplementary Fig. 1), and it was comparable to monosynaptic conductance between pvBCs (average synaptic conductance in BC-BC three pairs; 1.04 nS, 1.68 nS and 1.84 nS). Synaptic GABA_A_R- mediated peak conductance was 3.10 nS, 1.42-3.94 nS (median, quartiles) (P = 0.128, MW-test compared to G_aut_ in humans). The monosynaptic cell pairs are listed in Supplementary Table 1 with details on their spike kinetics and pv immunoreactivity.

**Supplementary figure 1.**
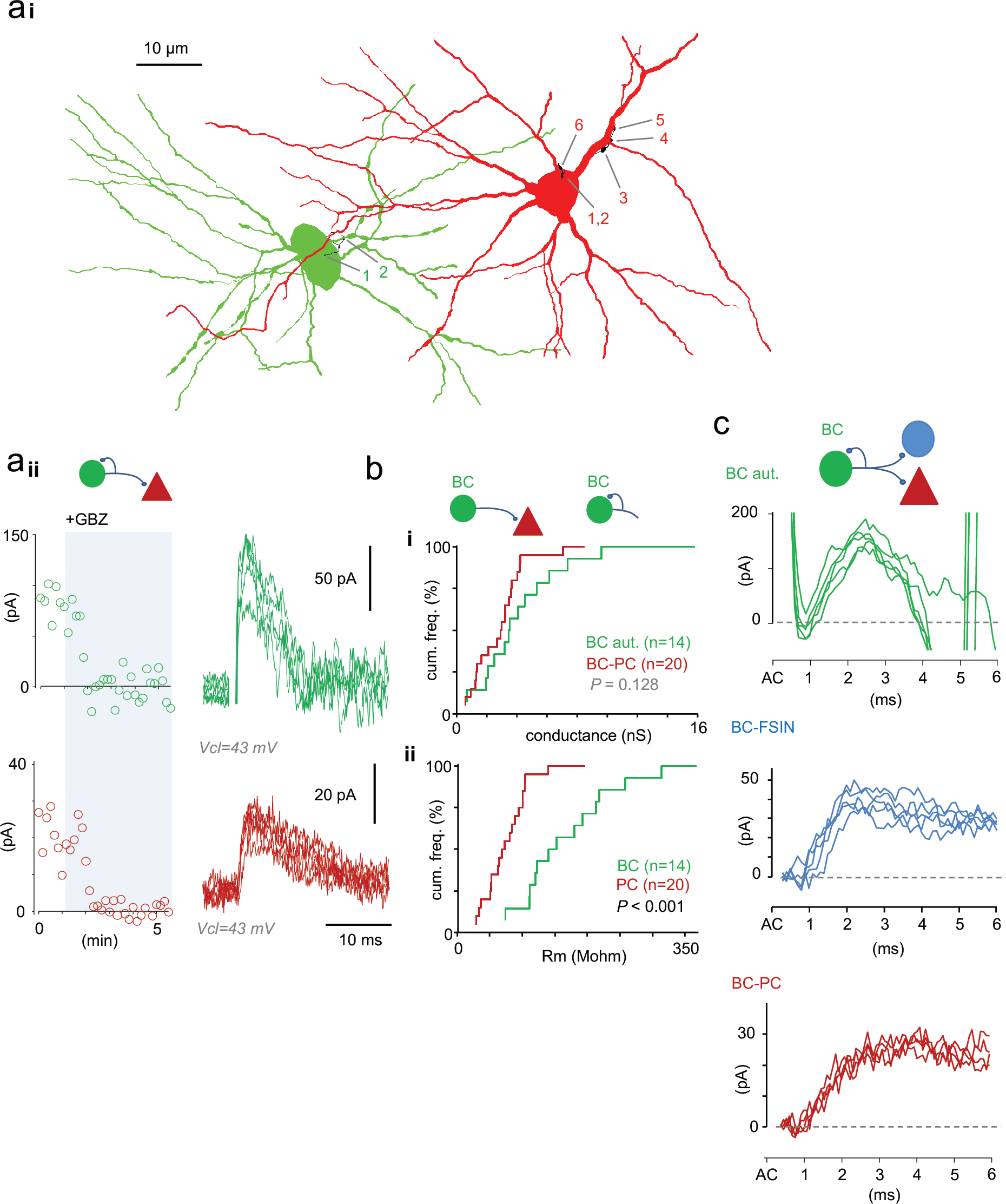
GABAergic self-innervation is comparable to synaptic connection from pvBC to L2/3 neurons. **a)** A paired recording of a synaptically connected pvBC (green)-to-PC (red) pair. (i) Close apposition pvBC axon boutons (black) showing autaptic and synaptic contacts. (ii) *Left:* plot shows autaptic (green symbols) and synaptic (red symbols) currents measured simultaneously in a paired recording. After a baseline, wash-in of gabazine (GBZ, 10 μM) blocks both currents. *Right:* superimposed individual traces of the autaptic and synaptic currents at baseline. **b)** Average autaptic GABA_A_R-mediated peak conductance is comparable to synaptic conductance from pvBC to PC. Autapses and synapses were recorded in separate experiments (GBZ not washed in 15 of 20 synaptic pair recordings). (i) On average, pvBC self-innervation (green, 14 pvBCs) shows similar peak conductance as monosynaptic IPSCs in L2/3 PCs (red, 20 pvBC-PC synaptic pairs). The autapse data are the same as those presented in Figure 2. (ii) Input resistance (R_m_ in resting condition) is higher in pvBCs (green) than in PCs (red) (P = 0.001, MW-test). This result suggests a theoretically stronger average shunting inhibition effect by autapses in pvBCs than by synapses in PCs. **c)** Triple recording from a pvBC with autapse and synaptic connection to a PC and to a FSIN. Note the similar rise kinetics of the autaptic current and IPSC in FSIN.

Because autaptic function has not been studied in nonhuman supragranular layer interneurons, we investigated L2/3 pvBCs in the mouse somatosensory cortex. We utilized mice expressing td-Tomato fluorophore preferably in parvalbumin GABAergic neurons (Chattopadhyaya B et al. 2004), confirmed each studied cell as fast-spiking (Supplementary Table 1), and visualized cells with fluorophore-streptavidin after filling with biocytin. Three cells were further selected for immunohistochemical investigation and confirmed to be immunopositive for pv, as illustrated in Fig. 2b_i_. We found GBZ-sensitive autapses in 11 of 19 mouse pvBCs, representing 57.9 % of the cells studied (Fig. 2b_ii,iii_). Figure 2b_i_ illustrates a visualized sample mouse pvBC with its axon forming close appositions close to its own perisomatic area. Mouse pvBCs (n = 11) had G_aut_ peaks of 4.09 nS, 1.93-5.67 nS, akin to what we discovered in human pvBCs (P = 0.72, MW-test) (Fig. 2c_i_). Autaptic current decay was defined in cells showing a large autaptic current (allowing confident monoexponential curve fitting r2 range 0.85 to 0.97) and a 3.43 ± 0.24 ms time constant (SW P = 0.091, n = 7).

Mouse pvBC input resistance under resting conditions (139.60 ± 6.69 MΩ, SW P = 0.566) was not significantly different from R_m_ in human pvBCs (183.83 ± 22.34 MΩ, SW P = 0.247, n = 14) (Welch’s t-test P = 0.077), but compared to mouse pvBCs, human pvBCs showed a wider value range (K-S test D = 0.500, P = 0.060) (Fig. 2c_ii_). On average, human and mouse pvBCs exhibited comparable G_aut+leak_ / G_leak_-values (SW P = 0.985, Student’s t-test P = 0.358 vs. human). This value shows the peak G_aut_ effect on total cell conductance and reflects its capacity to reduce cell excitability. (G_leak_ = membrane leak conductance in resting conditions; G_aut+leak_ = total cell input conductance during peak G_aut_). The G_aut_ relation to G_leak_ is illustrated in Fig. 2c_iii_.

### Temporal window for autaptic shunting inhibition in pvBCs following a spike

The inhibitory efficacy of GABA_A_R-mediated self-inhibition depends on G_aut_ and is related to the cell R_m_ and the relative proportion of G_aut_ and non-GABAergic spike afterhyperpolarization conductance (G_ahp_). We next investigated the relative strength of G_aut_ and G_ahp_ in pvBCs. The average G_ahp_ and G_aut_ traces fitted to monoexponential decay curves in one cell are illustrated in Fig. 3a_i_.

**Figure 3.**
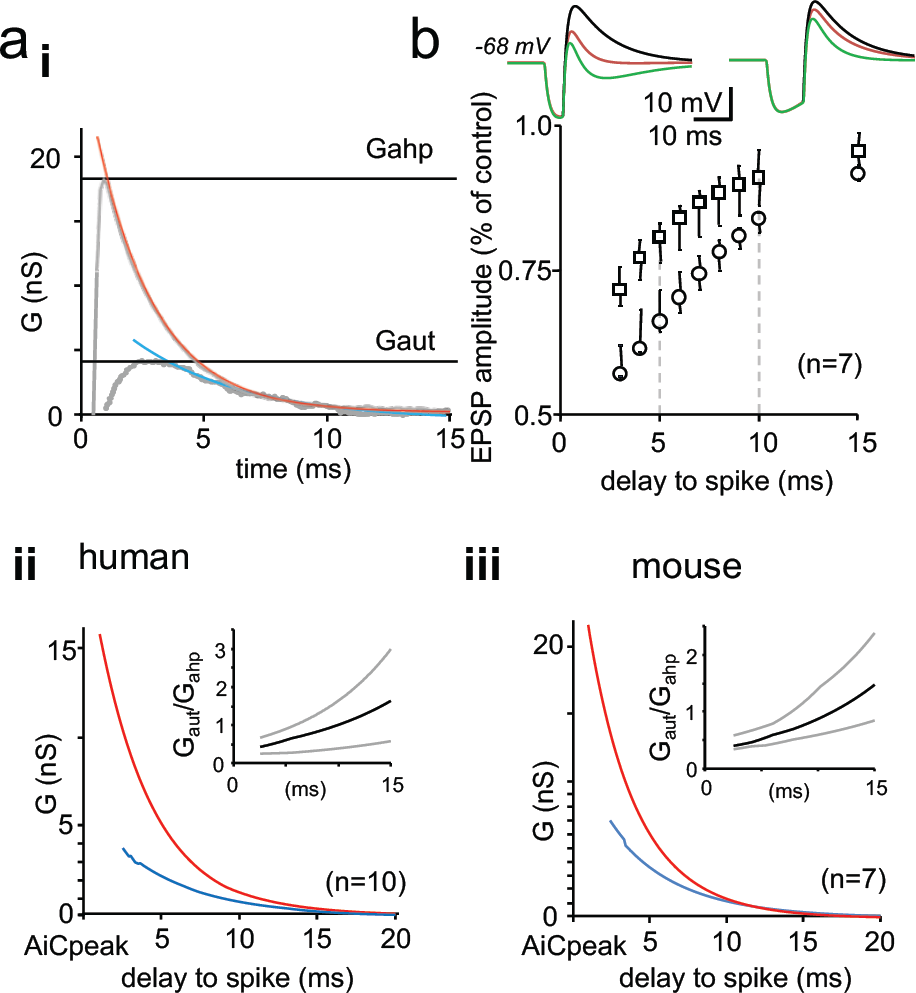
Time window for autaptic self-inhibition in pvBCs following a spike. **a)** Autaptic GABA_A_R-mediated conductance (G_aut_) overlaps with AHP current conductance (G_ahp_), increasing inhibition during AHP decay. (i) The two conductances G_ahp_ and G_aut_ in a human pvBC (gray lines, averages of 6 both). Horizontal lines indicate peak amplitudes. Red and blues curves show their monoexponential decay from the peak. Abscissa 0-time point indicates the pvBC spike inward current peak. (ii) Average monoexponential decay for G_ahp_ (red) and G_aut_ (blue) in ten human pvBCs showing the strongest autaptic component that demonstrates the temporal behavior of the two inhibitory conductances. Abscissa indicates a delay (in ms) to the spike inward current peak. The inset plot shows the relative proportion of G_aut_/G_ahp_ in the cells (median and quartiles). (iii) Corresponding conductances in mouse pvBCs. Inset shows the relative proportion of G_aut_ and G_ahp_. **b)** Relative contribution of G_aut_ is strongest during G_ahp_ decay. Simulation shows inhibition of EPSP in human pvBCs by G_aut_ after a spike. Simulation with a single-cell perisomatic model uses parameters recorded in individual human pvBCs (Supplementary Table 2). Simulated EPSP occurs with increasing delay to an action potential. Circle symbols show EPSP amplitude (median and quartiles) inhibited by G_ahp_ and G_aut_ after a spike. Box symbols show the EPSPs inhibited by G_ahp_ only in the absence of G_aut_. EPSP in each time point is normalized to an amplitude in the absence of both G_ahp_ and G_aut_. Simulations are from 6 pvBCs showing the most robust autaptic component (G_aut_ > 4.5 nS). Traces on top show EPSPs in one pvBC at 5 ms and 10 ms delay time points (green, in the presence of G_ahp_ and G_aut_; red, in the presence of G_ahp_ but not G_aut_; black, EPSP in the absence of G_ahp_ and G_aut_).

Human pvBCs showed a peak G_ahp_ of 17.35 ± 1.73 nS (SWP = 0.613, n = 14) measured from the action potential escape current outward component in the presence of GBZ (peak at 0.81 ± 0.07 ms delay to the inward action current peak, SW P = 0.096). The variability in the G_ahp_ peak between individual pvBCs probably reflects the cell soma size since the peak G_ahp_ value correlated with the cell capacitance (C_m_ = 32.42, 23.16-46.42 pF, n = 14) (Spearman’s r = 0.709, P = 0.019, n = 14) and G_leak_ (Pearson’s r = 0.664, P = 0.036, n = 14). However, none of the three parameters correlated with peak G_aut_. The AHP outward current decay time constant was 5.05 ± 0.51 ms (SW P = 0.14, n = 14) (Fig. 3a_1_). The average monoexponential decay curves for human pvBC G_ahp_ and G_aut_ are illustrated in Fig. 3a_ii_ for 10 cells with the most robust G_aut_ to demonstrate the temporal features.

The mouse G_ahp_ peak was 24.44 ± 2.49 nS (SW P = 0.076, n = 11), showing a 0.62 ± 0.059 ms (SW P = 0.103, n = 11) delay to the axon inward current. The AHP current had a decay time constant of 3.16 ± 0.25 ms (SW P = 0.061, n = 11). Thus, the G_ahp_ peak in mouse cells was higher than that in human pvBCs (P = 0.025 student’s t-test), and the AHP current in mice was moderately shorter than that in humans (decay tau average 3.16 vs. 5.05, P = 0.010, Student’s t-test). Fig. 3a_iii_ shows the average G_aut_ and G_ahp_ monoexponential decay in mouse cells with the most robust G_aut_ (n = 7).

Although G_ahp_ was larger than G_aut_ in human and mouse pvBCs, the role of GABA_A_R- mediated self-inhibition conductance became larger relative to G_ahp_ by increasing the temporal distance from the action potential, as demonstrated in the insets of Figs. 3a_ii,iii_.

To study the time window for efficient autaptic inhibition in human pvBCs, we utilized a computational single-cell perisomatic model using parameters measured in the cells above (including G_leak_, C_m_, and G_ahp_ and G_aut_ with peak times and decay times) (Connelly WM and G Lees 2010).

We simulated EPSPs elicited by EPSCs with features recorded in pvBCs (rise tau 0.2 ms, decay tau 1.2 ms, conductance 10 nS) (Szegedi V et al. 2016) at specific time points after pvBC action potential in six human pvBCs showing the most robust G_aut_. First, we ran simulations in the presence of G_aut_ and G_ahp_ as measured in individual cells. Next, we reproduced EPSPs in the presence of G_aut_ but in the absence of G_ahp_. In all simulations, the GABA_A_ reversal potential was set at the membrane potential of EPSP onset (Supplementary Table 2). Finally, as a reference, we ran EPSP simulations in the absence of both G_aut_ and G_ahp_. The membrane potential in the simulations was set to the value matching with control conditions.

The simulations demonstrated an effective postspike time window for autaptic inhibition during the first 10 ms delay from spike. The inhibitory effect of G_aut_ and G_ahp_ on the EPSP amplitude in the pvBCs is illustrated in Fig. 3b.

### Dynamic clamp reveals efficient GABA_A_R-mediated self-inhibition in human pvBCs

Next, we used whole cell dynamic clamp to generate somatic EPSPs in human L2/3 interneurons to study GABA_A_R-mediated self-inhibition in pvBCs. Dynamic clamp setting is well suited to investigate autaptic inhibition because it allows generation of EPSPs in pvBCs without possible disynaptic inhibition from neighboring interneurons (Molnar G et al. 2008; Komlosi G et al. 2012; Szegedi V et al. 2016).

We applied EPSCs with conductance and kinetic features measured in pvBCs (amplitude 1.5-5 nS, decay time constant 1.25 ms) (n =7 cells, resting membrane potential from −63 to −78 mV). In addition to a suprathreshold EPSP, we evoked two subthreshold EPSPs (2 – 9 mV), one preceding 40 ms the spike (prespike EPSP) and another time-locked to elicited action potential with a 5 ms delay (postspike EPSP). Following baseline, wash-in of GBZ (10 μM) selectively increased the postspike EPSP amplitude in pvBCs to 119.6 ± 3.3 % from baseline (mean of means ± sem, P = 0.0001, Student’s t-test, n = 7 cells) (SW P of data points in individual experiments from 0.127 to 0.995) that had no effect on prespike EPSP amplitude (102.0 ± 1.6 % of baseline, n = 7). The GBZ effect in pvBCs is illustrated in Fig. 4a_i-iii_. Similar experiments in nonFSINs (action inward current width 0.850 ± 0.041 ms vs. 0.600 ± 0.031 in pvBCs, P = 000193, t-test) failed to increase the prespike EPSP (amplitude 104.0 ± 1.5 % of baseline) or postspike EPSP (97.1 ± 4.0 % of baseline) (n = 8). GBZ wash-in (5 min) in nonFSINs is shown in Fig. 4b_i-iii_. The results confirm effective GABA_A_R-mediated autaptic inhibition in human pvBCs.

**Figure 4.**
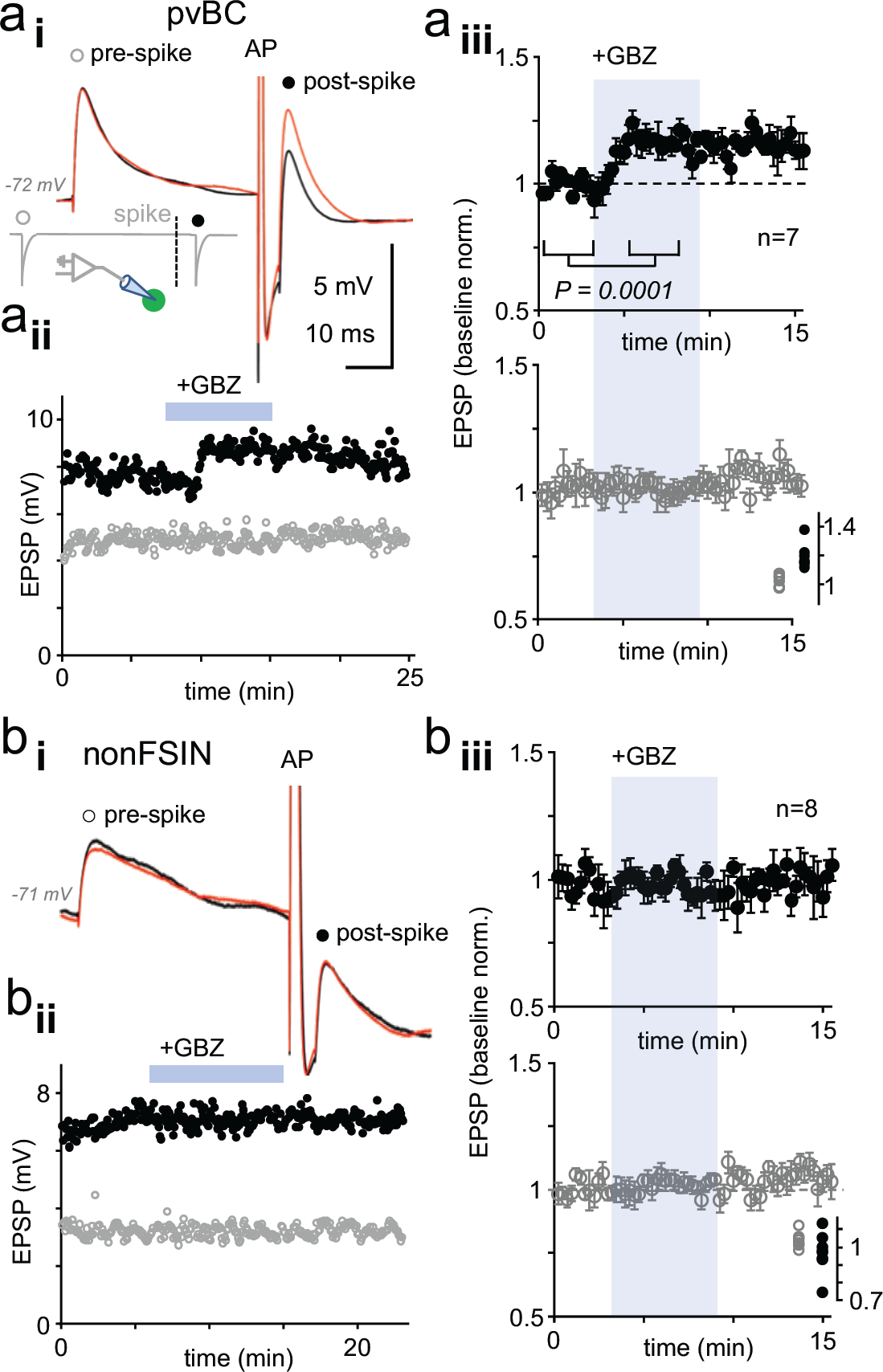
Dynamic clamp reveals autaptic inhibition of EPSPs in human pvBCs but not in nonFSINs. **a)** GABA_A_R-mediated self-inhibition of EPSP in pvBCs. A dynamic clamp experiment with two subthreshold EPSPs generated 40 ms before (prespike) and 5 ms after (postspike) a spike in pvBCs. Spike was evoked by suprathreshold EPSP. (i) Sample EPSPs (average of 6) in baseline conditions (black) and after wash-in of GBZ (10 μM) (red). Note the selective increase of the post-EPSP amplitude by GBZ. Inset schematic shows experimental design with prespike (open symbol), postspike (solid symbol) and suprathreshold (middle) inward conductance commands. The initiation of postspike EPSC conductance was time-locked to the peak of action potential evoked by the large EPSP. (ii) Amplitude of the pre-(open symbols) and postspike EPSPs (solid symbols) in the same experiment (amplitude from onset to peak). GBZ wash-in is indicated by a horizontal bar. (iii) Mean ± sem of 7 pvBCs (baseline-normalized). The shaded area indicates the wash-in of GBZ. Inset summarizes the baseline-normalized pre- and postspike amplitudes in the presence of GBZ in individual experiments. **b)** NonFSINs fail to show GABA_A_R-mediated self-inhibition of somatic EPSP. (i) Traces illustrate pre- and postspike EPSP (average of 6) in a nonFSIN during baseline (black) and in GBZ (10 μM) (red). (ii) Amplitude of the pre- and postspike EPSPs plotted in the same experiment. (iii) Mean ± sem of 8 nonFSINs. Inset summarizes postspike EPSP amplitudes in GBZ (baseline-normalized).

### Autapses inhibit repetitive spike firing in human pvBCs

Finally, we used dynamic clamp to test whether somatic self-inhibition is sufficient to control pvBC action potential firing. By applying suprathreshold excitatory inward current conductance (8 – 21 nS, decay time constant 4-5 ms), we elicited firing of action potential doublets in fast-spiking pvBCs in the presence of GBZ (spike interval 6.06 ms, 5.62-6.58 ms, n = 198 doublets, 3 cells). First, we found that wash-in of GBZ shortened the spike doublet interval (Fig. 5 ai_-ii_), indicating GABA_A_R-mediated inhibition of the 2^nd^ spike initiation in baseline conditions. In the continuous presence of applied GBZ, in addition to the spike doublet-triggering EPSC conductance, inhibitory conductance with onset delay (1 ms to 1st spike), amplitude (1-10 nS) and decay time (5 ms) akin to autaptic GABA_A_R-mediated responses were generated in dynamic clamp in pvBCs (see Methods) (Fig. 5b_i_). We found that by the autaptic inhibitory conductance, the doublet spike interval was elongated (Fig. 5b_ii_) or the 2^nd^ spike probability was significantly reduced (Figs. 5b_ii_, c-d) (ANOVA on ranks with Dunn’s post hoc test, Wilcoxon signed rank test in Fig. 5c). Applying 2.5 nS-10 nS autaptic inhibition by dynamic clamp reduced the 2nd spike probability in the three experiments to 0.34, 0.02-0.67 from 0.93, 0.69-1.00 in 0 nS autapse conditions (*P* = 0.048, MW test). Furthermore, as low as 1 nS autapse (tested in two of the cells) was capable of reducing 2nd spike initiation (Fig. 5c). Thus, the results demonstrate that autaptic conductance measured in human pvBCs is sufficient to control spiking of the cells.

**Figure 5.**
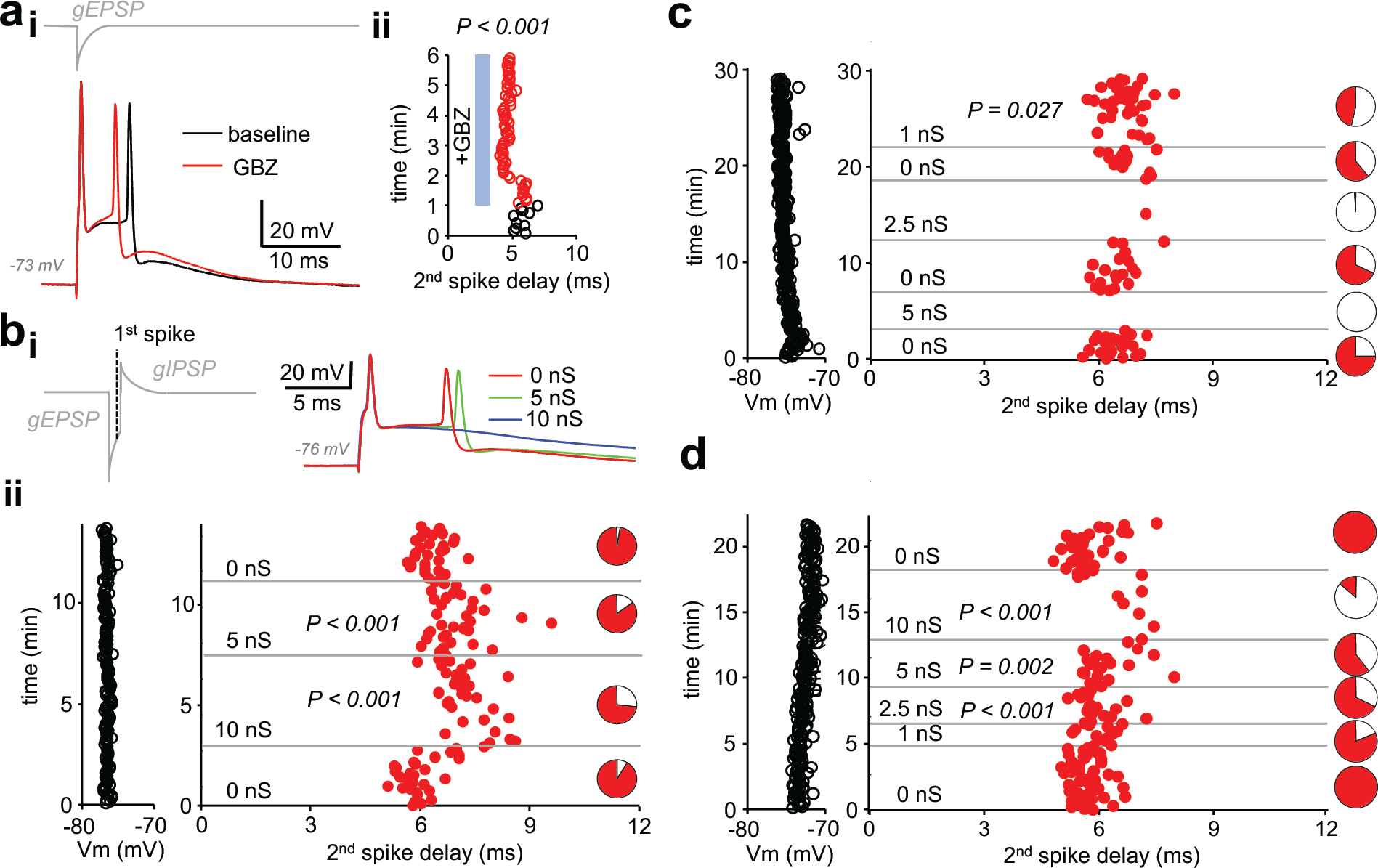
Autaptic inhibitory conductance regulates firing in human pvBCs. **a)** GBZ shortens a time delay between two spikes evoked by EPSP in dynamic clamp. (i) Sample traces in a pvBC evoked by EPSP in dynamic clamp during baseline and after wash-in of GBZ (μM). Trace on top (gray) shows EPSP conductance command (gEPSP). (ii) Delay from the 1^st^ to the 2^nd^ spike plotted in the same experiment at baseline (black line circles) and during wash-in of GBZ (red line circles). **b)** Inhibition of doublet spiking by natural autaptic conductance demonstrated with dynamic clamp in human pvBCs. Experiments in the presence of GBZ show decreased probability and increased delay of the second spike when autaptic conductance is introduced with dynamic clamp. Spikes are evoked by large single EPSP as in (a). (i) *Left:* schematic traces illustrate dynamic clamp command schematic with EPSP conductance followed by GABA_A_R autaptic IPSP conductance (with 1 ms onset delay to 1^st^ spike generated by EPSP). *Right:* traces showing the evoked spikes. (ii) *Left:* pvBC membrane potential during an experiment. *Middle:* red dots indicate the second spike onset delay to the first spike elicited by dynamic clamp EPSP (interval 5 s) when it was followed by autaptic IPSC conductance (5 nS or 10 nS). *P*-values show significant spike delay increased by autaptic conductance compared to baseline (ANOVA on ranks, post hoc Dunn’s pairwise test against all 0 nS data as control). *Right:* pie charts show the 2nd spike probability (red area) in all cycles with the specific IPSC conductance. Note the increased failure rate (white pie chart area) with 5 nS or 10 nS inhibitory autapse. **c-d)** Similar experiments as in (b) in two additional pvBCs in which autaptic inhibition (1 nS −10 nS) was more clearly seen in the probability of the 2nd spike (white area in pie charts). Wilcoxon signed rank test in (c), ANOVA on ranks with post hoc Dunn’s pairwise test in (d).

## DISCUSSION

Although autaptic self-inhibitory connections have been reported in GABAergic interneurons in rodents and some other experimental animals, their existence and function in identified interneurons in the human brain have remained virtually unknown. Furthermore, studies on autapses in animal experiments have heavily focused on neurons in infragranular layers of the neocortex, while the function of autapses in the supragranular layers has remained unexplored (Tamas G et al. 1997; Bacci A et al. 2003; Bacci A and JR Huguenard 2006; Connelly WM and G Lees 2010; Jiang M et al. 2012; Jiang M et al. 2015). Therefore, it has been unknown whether robust autaptic self-inhibition is a general feature in the neocortex operating in various neocortical layers and different species or whether strong GABAergic self-inhibition is a specialization in deep neocortical interneurons reported in rodents. Our study shows that GABA_A_R-mediated self-inhibition is a regular feature of supragranular layer pvBCs in humans and mice. pvBC axon terminals self-innervate their soma and proximal dendrites, and electron microscopic investigation shows that these contacts form ‘autaptic densities’ in the interneurons. The regular occurrence of autapses in pvBCs in different layers and areas of the neocortex, their robust inhibitory efficacy in pvBCs and their rare occurrence in nonFSINs indicate that self-inhibition is a common but cell type-specific microcircuit feature in the mammalian neocortex.

### Robust perisomatic self-inhibition regulates pvBC excitability and firing in distinct cortical layers and in different mammalian species

Here, our results in the supragranular layer together with other studies in the infragranular layers show that autapses efficiently control excitability in pvBCs after spikes (Bacci A and JR Huguenard 2006; Connelly WM and G Lees 2010; Jiang M et al. 2012; Deleuze CB, G.; Pazienti, A.; Mailhes, C.; Aguirre, A.; Beato, M.; Bacci, A. 2018). Autapses are present in similar proportions and show comparable inhibitory efficacy in pvBCs of superficial and deep neocortical layers (Bacci A and JR Huguenard 2006). In addition, our results here with human and mouse cells confirm that autapses have similar occurrence and strength in pvBCs in mice, rats and humans. Correspondingly, self-innervation is rare in human nonFSINs, similar to rodent nonFSINs (Bacci A et al. 2003; Jiang M et al. 2012). In general, GABAergic self-inhibition conductance in pvBCs is comparable to the synaptic inhibition these interneurons exert on neighboring layer 2/3 neurons.

Autaptic terminals on pvBC are heavily perisomatic, whereas nonFSINs, when autapses are found in them, self-innervate their own dendrites (Tamas G et al. 1997) akin to synaptic contacts made by these interneurons to other neurons (Favuzzi E et al. 2019). Information on the self-innervation subcellular target motifs as well as knowledge on autaptic inhibition strength in identified neurons is critical for understanding the operation of autaptic microcircuits in individual neurons as well as in a neuronal network.

In pvBCs, perisomatic GABAergic self-inhibition overlaps with much stronger AHP potassium conductance. However, the autaptic conductance onset delay and late peak time indicate that GABA_A_R-mediated self-inhibition is most influential during AHP conductance decay. In this way, the autaptic activity prolongs the time window for somatic inhibitory conductance up to 10 ms following a spike. We demonstrated this finding using a single-cell model as well as dynamic clamp experiments and showed that autaptic inhibition elongates the action potential interval (Bacci A and JR Huguenard 2006; Connelly WM 2014; Guo D et al. 2016; Yilmaz E et al. 2016; Deleuze CB, G.; Pazienti, A.; Mailhes, C.; Aguirre, A.; Beato, M.; Bacci, A. 2018) through shunting inhibition in human pvBCs where the GABA_A_ reversal potential is close to the resting membrane potential (Verheugen JA et al. 1999; Connelly WM and G Lees 2010). Autaptic self-innervation strength shows substantial variability between individual pvBCs, and it is possible that autaptic GABAergic contacts undergo strength-regulating plasticity (Castillo PE et al. 2011; Donato F et al. 2013; Griffen TC and A Maffei 2014; Lourenco J et al. 2014; Lourenço JMDS, A.; Deleuze, C.; Bigot, M.; Pazienti, A.; Aguirre, A.; Giugliano, M.; Ostojic, S.; Bacci A. 2019).

Overall, perisomatic self-innervation in human and mouse pvBCs reinforces self-inhibition after a spike. This process adjusts the pvBC firing interval and rhythmic inhibition from pvBCs to other neurons (Guo D et al. 2016; Yilmaz E et al. 2016; Deleuze CB, G.; Pazienti, A.; Mailhes, C.; Aguirre, A.; Beato, M.; Bacci, A. 2018).

### pvBC features in humans and mice

Comparison of autapses or cell input resistance showed no difference between human and mouse pvBCs. However, close investigation of the data reveals that compared to mouse pvBCs, human pvBCs exhibit a wide range of autapse conductance as well as cell input resistance. One potential explanation for this finding is human tissue material diversity. Thus, cortical region specificity and patient gender and age may partly be behind the parameter variability in human pvBCs.

However, among various cellular features characterized in humans but not in rodent neocortex (Blazquez-Llorca L et al. 2010; Testa-Silva G et al. 2010; Molnar G et al. 2016; Szegedi V et al. 2016; Wang B et al. 2016; Beaulieu-Laroche L et al. 2018; Boldog E et al. 2018; Goriounova NA et al. 2018; Kalmbach BE et al. 2018; Pruunsild P and H Bading 2019), human neocortical neurons exhibit higher R_m_ than their rodent counterparts do (Eyal G et al. 2016; Poorthuis RB et al. 2018). Although our data here showed that human and mouse pvBC input resistance values were not different on average (showing p value of 0.10), human neurons had individual cells showing clearly higher R_m_ than did those found in any of the mouse pvBCs (Poorthuis RB et al. 2018).

In line with this result, we found that AHP conductance was generally higher in mouse cells than in human cells since high R_m_ requires less current to generate an equal amplitude potential. Indeed, human pvBC showed a correlation between AHP conductance and membrane leak conductance. Thus, high R_m_ pvBCs, particularly in the human neocortex, need smaller conductance for efficient inhibition than do pvBCs in mice.

## Conclusions

Firing of pvBCs coordinates synchrony of neuronal networks in the cortex (Hu H and P Jonas 2014; Lu J et al. 2017; Cardin JA 2018). Through robust autapses, pvBCs adjust their temporal firing interval and correspondingly set inhibition in their target neurons. This mechanism may be essential in setting cell assembly discharges in the neocortex (Molnar G et al. 2008; Toth K et al. 2018; de la Prida LM and G Huberfeld 2019) during associative memory processing (Kucewicz MT et al. 2014) and memory retrieval (Vaz AP et al. 2019). In addition, pvBC self-inhibition may contribute to pyramidal cell disinhibition during the induction of L2/3 LTP associated with learning (Williams LE and A Holtmaat 2019).

## METHODS

### Ethics statement

All procedures were performed according to the Declaration of Helsinki with the approval of the University of Szeged Ethical Committee and Regional Human Investigation Review Board (ref. 75/2014). For all human tissue material, written consent was obtained from patients prior to surgery. Tissue obtained from underage patients was provided with agreement from a parent or guardian. In 5 of the 20 pvBC-PC cell pairs reporting IPSC conductance, some other data parameters (excluding conductance reported here) have been reported in a previous manuscript (Szegedi V et al. 2017).

### Human brain slices

Neocortical slices were sectioned from material that had to be removed to gain access for the surgical treatment of deep-brain targets from the left and right frontal, temporal or occipital areas. In some cases, tissue from neocortical operations was used when it was nonpathological. In these latter cases, small pieces of nonpathological tissue had to be removed in the surgery to obtain access to pathological targets in the folded neocortex. The patients were 10-85 years of age, and samples from males and females from either the left or right hemisphere were included. Details including patient gender, age, resected neocortical area and pathological target diagnosis are reported for all tissue samples used in this study in Supplementary Table 1. Anesthesia was induced with intravenous midazolam and fentanyl (0.03 mg/kg, 1–2 lg/kg, respectively). A bolus dose of propofol (1–2 mg/kg) was administered intravenously. The patients received 0.5 mg/kg rocuronium to facilitate endotracheal intubation. The trachea was intubated, and the patient was ventilated with an O_2_/N_2_O mixture at a ratio of 1:2. Anesthesia was maintained with sevoflurane at a care volume of 1.2–1.5. Following surgical removal, the resected tissue blocks were immediately immersed into a glass container filled with ice-cold solution in the operating theatre. The solution contained (in mM): 130 NaCl, 3.5 KCl, 1 NaH_2_PO_4_, 24 NaHCO_3_, 1 CaCl_2_, 3 MgSO_4_, 10 D(+)-glucose and was saturated with 95 % O_2_/5 % CO_2_. The container was placed on ice in a thermally isolated transportation box where the liquid was continuously gassed with 95 % O_2_/5 % CO_2_. Then, the tissue was transported from the operating theatre to the electrophysiology lab (door-to-door in maximum 20 min), where slices of 350 μm thickness were immediately prepared from the block with a vibrating blade microtome (Microm HM 650 V). The slices were incubated at 22-24°C for 1 h, when the slicing solution was gradually replaced by a pump (6 ml/min) with the solution used for storage (180 ml). The storage solution was identical to the slicing solution, except for 3 mM CaCl_2_ and 1.5 mM MgSO_4_.

### Pv+ cells in mouse brain slices

Transversal slices (350 μm) from somatosensory cortex were prepared (Kotzadimitriou D et al. 2018) from 4- to 6-week-old heterozygous male CB6-Tg(Gad1-EGFP)G42Zjh/J-mice (The Jackson Laboratory, stock 007677, GAD67-GFP G42 line) expressing td-Tomato fluorophore preferably in parvalbumin GABAergic neurons (Chattopadhyaya B et al. 2004). Cells were confirmed to be fast-spiking, showing fast spike kinetics and a high-frequency non-accommodation firing pattern for suprathreshold depolarizing 500 ms pulses (Supplementary Table 1). Cells were visualized with streptavidin Alexa488 (1:2000, Jackson ImmunoResearch Lab, Inc.) and analyzed by eye under epifluorescence microscopy to exclude axo-axonic cells. Three cells were selected for pv immunoreactivity, and they were all immunopositive for pv.

### Electrophysiology

Recordings were performed in a submerged chamber (perfused 8 ml/min) at 36–37°C. Cells were patched using a water-immersion 20x objective with additional zoom (up to 4x) and infrared differential interference contrast video microscopy. Micropipettes (5–8 MΩ) for whole-cell patch-clamp recording were filled with intracellular solution with physiological or elevated intracellular chloride [Cl^−^]i. The content of the solution for voltage clamp recordings with physiological [Cl^−^]i was (in mM): 126 K-gluconate, 8 KCl, 4 ATP-Mg, 0.3 Na2–GTP, 10 HEPES, and 10 phosphocreatine (pH 7.20; 300 mOsm) with 0.3 % (w/v) biocytin. Current clamp recordings with elevated [Cl^−^]i contained 130 mM KCl instead. Recordings were performed with a Multiclamp 2B amplifier (Axon Instruments) and low-pass filtered at 6-8 kHz cut-off frequency (Bessel filter). Series resistance (Rs) and pipette capacitance were compensated in current clamp mode and pipette capacitance in voltage clamp mode. Rs was monitored and recorded continuously during the experiments. Voltage clamp recordings were discarded if the Rs was higher than 25 MΩ or changed by more than 20 %. Liquid junction potential error was corrected in all membrane potential values. The access resistance of the recording electrode was measured, and its effect on the clamping potential error was corrected in nominal somatic potential reading. Resting membrane potential was recorded 1-3 min after break-in to whole cell. Cell capacitance and input resistance were measured in current clamp using -50-100 pA, 600 ms steps delivered at resting membrane potential.

Single-spike firing in current clamp or in voltage clamp was induced by a 50 ms depolarizing suprathreshold step from the resting membrane potential. The autaptic GBZ-sensitive outward current amplitude was 150.2 pA, 60.0 to 177.5 pA (median, quartiles, n = 14). Single-spike firing in AACs was elicited with 1-2 ms suprathreshold depolarization. Synaptic IPSCs in pvBC-PC pairs were recorded at steady postsynaptic −43 mV to −55 mV clamping potential. Conductance was calculated from averaged (at least 12 events) current peak amplitude and the GABA_A_ current driving force for each cell by Ohm’s law formula. The GABA_A_ current reversal potential with the recording solution containing physiological Cl^−^ was −73 mV. The reversal potential for AHP was −92 mV. Evoked autaptic and synaptic responses were analyzed using Clampfit or Spike2 programs as described in (Szegedi V et al. 2017). Cell input resistance and capacitance were measured in the current clamp at the resting membrane potential.

### Dynamic clamp

To emulate the EPSPs and IPSCs in basket cells and other interneurons, a software-based dynamic clamp system was employed. Current injections were calculated by the StdpC2017 software (Kemenes I et al. 2011) through a MIO-16E-4 analog/digital card (National Instruments Inc., Hungary) based on voltage signals of the electrode. We ran the dynamic clamp on a separate computer from our experimental data acquisition system (recording cell membrane potential) to record and verify dynamic clamp output (conductance and EPSCs). Sub- and suprathreshold EPSCs were evoked using a decay time constant of 1.25 ms and a reversal potential of 0 mV. The peak conductance (1.5 – 8 nS) for subthreshold EPSP was set to evoke a 2-9 mV peak amplitude response, and the onset of postspike EPSC was triggered by a preceding spike being time locked to it with a 5 ms onset delay. EPSCs triggering the spike had peak conductance up to 20 nS. Autaptic IPSCs had 1 nS-10 nS conductance, a decay time constant of 5 ms and a reversal potential of −78 mV. The onset of IPSC was triggered by a preceding spike with a 1 ms delay.

### Single cell model

In the simulation of somatic EPSP inhibition by autaptic shunting, real experimental data from individual basket cells were used. Simulation of a basket cell membrane potential was performed using a NEURON 7.6.5 simulator (Carnevale NT HMC, UK: Cambridge University Press 2006). The membrane capacitance was 1 μF/cm2, and the size of the soma was determined so that the total cell capacitance matched the actual measured value in each cell. Conductance of leak current was retrieved from experimental data. The reversal potential of the leak current determining the resting membrane potential was set to −68 mV. We defined 3 point processes using a current input model to simulate AHP, autapse and incoming EPSP (Schmidt-Hieber C et al. 2007). For AHP conductance, the rise tau was set to 0.1 ms, and the reversal potential was set to −90 mV. Decay tau and conductance maximum amplitude were derived from experimental data for each cell (see Supplementary Table 2). The GABA_A_R current reversal potential was set to the resting membrane potential and peaked at 3 ms after AHP onset. Autaptic conductance maximum amplitude, rise and decay tau were obtained from the experimental data. EPSPs were modeled using rise tau 0.2 ms, decay tau 1.2 ms, conductance of 10 nS with reversal potential 0 mV. The EPSP onset delay was set to 3-15 ms from AHP current onset with varying steps (1 ms interval). Reference EPSP (without AHP or autapse) was evoked for each onset delay response at the Vm the EPSP onset showed in the presence of AHP and autapse. This reference was adjusted by setting the initial Vm to respective values in the model.

### Data analysis

Data were acquired with Clampex software (Axon Instruments) and digitized at 10–50 kHz. The data were analyzed off-line with pClamp (Axon Instruments), Spike2 (version 8.1, Cambridge Electronic Design), OriginPro (OriginLab Corporation) and IgorPro (WaveMetrics Inc.) and SigmaPlot software. Spike kinetics in the interneurons were measured as the axon potential inward current component width. Synaptic response parameters were analyzed as described previously (Szegedi V et al. 2016). Autaptic currents and potentials were defined by subtracting trace averages (of at least 6 events) in gabazine from traces evoked by preceding trials. Peak conductance was calculated from currents in voltage clamp according to Ohm’s law. The time-to-peak value was defined from the current onset to its maximal value.

### Statistics

Data are presented as the mean ± s.e.m, when showing n ≥ 7 with a parametric distribution. Normality was tested with the Shapiro-Wilk test (SW P value > 0.05). Otherwise, the data are shown as the median with interquartile range (of lower and upper quartile). Correspondingly, for statistical analysis, t-test, Mann-Whitney U-test (MW test), or Wilcoxon signed-rank test was used. In addition, the Kolmogorov–Smirnov test was used to test nonparametric probability distributions. Differences were accepted as significant if P < 0.05.

### Cell visualization

Biocytin-filled cells were visualized with either Alexa488-(1:500) or Cy3-streptavidin (1:400, Jackson ImmunoResearch Lab, Inc.) for anatomical and immunohistochemical investigation. After electrophysiological recording, slices were immediately fixed in a fixative containing 4 % paraformaldehyde and 15 % picric acid in 0.1 M phosphate buffer (PB, pH = 7.4) at 4°C for at least 12 h and then stored at 4°C in 0.1 M PB with 0.05 % sodium azide as a preservative. For some slices, 0.05 % glutaraldehyde was added before use for electron microscopy studies (Supplementary Table 1). All slices were embedded in 10 % gelatin and further sectioned into slices of 50 μm thickness in the cold PB using a vibratome VT1000S (Leica Microsystems). After sectioning, the slices were rinsed in 0.1 M PB (3 × 10 min) and cryoprotected in 10–20 % sucrose solution in 0.1 M PB. After this, the slices were frozen in liquid nitrogen and thawed in 0.1 M PB. Finally, they were incubated in fluorophore-conjugated streptavidin (1:400, Jackson ImmunoResearch Lab, Inc.) in 0.1 M Tris-buffered saline (TBS, pH 7.4) for 2.5 h (at 22–24°C). After washing with 0.1 M PB (3 × 10 min), the sections were covered in Vectashield mounting medium (Vector Laboratories Inc.), placed under cover slips, and examined under an epifluorescence microscope (Leica DM 5000 B).

### Cell reconstruction and anatomical analyses

Sections selected for immunohistochemistry and cell reconstruction were dismounted and processed (see Immunohistochemistry paragraph). Some sections for cell structure illustrations were further incubated in a solution of conjugated avidin-biotin horseradish peroxidase (ABC; 1:300; Vector Labs) in Tris-buffered saline (TBS, pH = 7.4) at 4°C overnight. The enzyme reaction was revealed by the glucose oxidase-DAB-nickel method using 3’3-diaminobenzidine tetrahydrochloride (0.05 %) as the chromogen and 0.01 % H_2_O_2_ as the oxidant. Sections were further treated with 1 % OsO_4_ in 0.1 M PB. After several washes in distilled water, sections were stained in 1 % uranyl acetate and dehydrated in an ascending series of ethanol concentrations. Sections were infiltrated with epoxy resin (Durcupan) overnight and embedded on glass slides. For the cells visualized in the figures, three-dimensional light microscopic reconstructions from one or two sections were carried out using the Neurolucida system with 100x objective (Olympus BX51, Olympus UPlanFI). Images were collapsed in the z-axis for illustration. The somatodendritic region in the 50 μm-thick section was studied for close appositions with filled axons traced back to the soma. Neurolucida explorer software was used to measure the distance of close appositions in dendrites to the soma in images visualized using the same computer software. The perisomatic area of (50 μm thick section) of nine DAB-visualized and identified axo-axonic cells was studied under a light microscope (100x magnification) for axonal contacts. The study included 5 biocytin-filled AACs in the data archive.

### Immunohistochemistry

Free-floating sections were washed 3 times in TBS-TX 0.3 % (15 min) at 22–24°C and then moved to 20 % blocking solution with horse serum in TBS-TX, 0.3 % for pv staining and 10 % blocking solution for vesicular GABA transporter (vgat) staining. The sections were incubated in primary antibodies diluted in 1 % serum in TBS-TX 0.3 % over three nights at 4°C, and placed in relevant fluorochrome-conjugated secondary antibodies in 1 % blocking serum in TBS-TX 0.3 % overnight at 4°C. Sections were first washed in TBS-TX 0.3 % (3 × 20 min) and later in 0.1 M PB (3 × 20 min) and mounted on glass slides with Vectashield mounting medium (Vector Lab, Inc.).

The characterizations of antibodies used in humans; (mouse anti-pv, 1:500, Swant, Switzerland, www.swant.com, clone: 235) and (rabbit anti-vgat, 1:500, Synaptic Systems, Germany, www.sysy.com, AB_887871). Fluorophore-labeled secondary antibodies were (DAM DyLight 488 donkey anti mouse, 1:400, Jackson ImmunoResearch Lab. Inc., www.jacksonimmuno.com) or (DAM Cy3 donkey anti mouse, 1:400, Jackson ImmunoResearch Lab. Inc., www.jacksonimmuno.com) and (DARb Cy5 donkey anti-rabbit, 1:500, Jackson ImmunoResearch Lab. Inc.). Antibodies in mice were (goat anti-pv, 1:2000, Swant, AB_10000343) and (DAGt Cy5 donkey anti-goat, 1:400, Jackson ImmunoResearch Lab. Inc.). The labeling of neurons by biocytin and immunoreactions was evaluated using first epifluorescence (Leica DM 5000 B) and then laser scanning confocal microscopy (Zeiss LSM880). All micrographs presented are confocal fluorescence images.

### Electron microscopy

Sections containing pvBC soma were re-embedded, and 65 nm thick ultrathin sections were cut with an ultramicrotome (RMC MT-XL). Ribbons of the sections were collected on Formvar-coated copper grids and examined with a JEOL JEM-1400Plus electron microscope. Images were taken by an 8 M pixel CCD camera (JEOL Ruby).

## ACKNOWLEDGMENTS

This work was supported by the National Brain Research (Nemzeti Agykutatási) program (KL, MP, VS, JB, GM and GT), the ERC INTERIMPACT project (GT), the Hungarian Academy of Sciences (GM, GT and VS), and University of Szeged Open Access Fund (Grant number: 4373). We acknowledge Leona Mezei for technical assistance.

---

**Supplementary table 1.** Details of recorded human neurons and information on the resected human cortical tissue. Columns from left to right show the figure in which the cell recording data are shown, cell filing code, indication for successful recovery of cell with streptavidin visualization (strept, NA indicates unsuccessful cell recovery), immunoreaction for parvalbumin (pv) and vesicular GABA transported (vGAT), action current inward component width (acw) and maximal firing frequency when tested (defined during first 100 ms of depolarizing pulse). pvBCs showing no evidence for autapses are shown in red. Details of resected tissue are shown in blue, showing patient gender, age, hemisphere, cortical area and diagnosed primary pathology. “no info” = ventriculostomy operations, and there is no info about the side and location in the original documentation.

|  | cell code | strept | immuno | acw | gender | age | hemisphere | neocortical area | diagnosed pathology |
| --- | --- | --- | --- | --- | --- | --- | --- | --- | --- |
| **Fig 1 exps** |  |  |  |  |  |  |  |  |  |
| **GBZ** | V041017_4 | strept | PV+, vGAT+ | 0.5 | female | 69 | right | occipital | subcortical neoplasia |
|  | V091117_1 | strept | Pv, vGAT? | 0.5 | female | 63 | right | temporal | hydrocephalus |
|  | V231117_1 | strept | vGAT+ | 0.7 | female | 68 | right | temporal | hydrocephalus |
|  | V071217_1 | strept | PV+ | 0.5 | female | 76 | right | frontal | hydrocephalus |
|  | V130618_2 |  |  | 0.5 | female | 35 | right | temporal | subcortical neoplasia |
| **BAPTA** | k190918_1 | NA |  | 0.4 | male | 21 | right | temporo-occipital | subcortical cystic neoplasm |
|  | v190918_4 | strept |  | 0.6 | male | 21 | right | temporo-occipital | subcortical cystic neoplasm |
|  | v190918_5 | strept |  | 0.5 | male | 21 | right | temporo-occipital | subcortical cystic neoplasm |
|  | v091018_1 | NA |  | 0.6 | female | 66 | left | frontal | cortical metaplasia |
|  | v171018_1 | NA |  | 0.6 | male | 33 | right | frontal | hydrocephalus |
|  | v171018_2 | NA |  | 0.5 | male | 33 | right | frontal | hydrocephalus |
| no aut | V041017_3 | strept | Pv+<br>vGAT + | 0.7 | female | 69 | right | occipital | subcortical neoplasia |
|  | v041017_5 | NA |  | 0.5 | female | 69 | right | occipital | subcortical neoplasia |
| **Fig 2 and 3**<br>**exps** |  |  |  |  |  |  |  |  |  |
| aut in exp | k151217_2 | strept | PV,<br>vGAT<br>non<br>concl | 0.65 | female | 19 | right | frontal | subcortical neoplasia |
|  | k080917_1 | NA |  | 0.5 | male | 45 | right | fronto-<br>temporal | subcortical neoplasia |
|  | k121217_1 | strept | PV+,<br>vGAT+ | 0.5 | female | 68 | right | temporal | hydrocephalus |
|  | k150218_1 | strept | PV non<br>concl | 0.7 | female | 67 | right | occipital | subcortical neoplasia |
|  | k151217_3 | strept | PV+,<br>vGAT+ | 0.6 | female | 19 | right | frontal | subcortical neoplasia |
|  | k150917_2 | strept | PV non<br>concl | 0.6 | female | 63 | right | occipital | subcortical neoplasia |
|  | k270218_1 | strept | PV non<br>concl | 0.65 | male | 34 | left | frontal | subcortical neoplasia |
|  | k260118_1 | strept | pv+ | 0.6 | female | 54 | right | frontal | hydrocephalus |
|  | k071217_1 | strept | PV+ | 0.6 | female | 76 | right | frontal | hydrocephalus |
|  | k280218_1 | strept | PV+ | 0.5 | female | 41 | right | frontal | hydrocephalus |
|  | k121217_3 | strept | pv+ | 0.5 | female | 68 | right | temporal | hydrocephalus |
|  | k231117_3 | strept | PV,<br>vGAT<br>non<br>concl | 0.65 | female | 68 | no info | no info | hydrocephalus |
|  | k260118_2 | strept | PV+ | 0.65 | female | 54 | right | frontal | hydrocephalus |
|  | k200218_6 | strept | PV+ | 0.45 | female | 54 | right | frontal | aneurism |
| no aut | k280218_1 | strept | PV+ | 0.35 | female | 41 | right | frontal | hydrocephalus |
|  | k250118_1 | strept | PV+ | 0.35 | female | 50 | left | occipital | astrocytoma |
|  | k250118_2 | strept | PV+ | 0.6 | female | 50 | left | occipital | astrocytoma |
|  | k200218_3 | strept | PV+ | 0.6 | male | 56 | right | occipital | hydrocephalus |
|  | k200218_4 | strept | PV+ | 0.45 | female | 54 | right | frontal | aneurism |
|  | k270218_2 | strept | PV non<br>concl. | 0.55 | male | 34 | left | frontal | subcortical neoplasia |
|  | k200218_5 | NA |  | 0.6 | female | 54 | right | frontal | aneurism |
|  | k231117_2 | strept | PV+ | 0.5 | female | 68 | no info | no info | hydrocephalus |
|  | k250118_3 | strept | PV+ | 0.5 | female | 50 | left | occipital | astrocytoma |
| **Suppl Fig 1**<br>**exps** |  |  |  |  |  |  |  |  |  |
|  | k231117_2 | strept | PV+ | 0.7 | female | 68 | right | temporal | hydrocephalus |
|  | k150218_1 | strept | PV non<br>concl | 0.65 | female | 67 | right | occipital | subcortical neoplasia |
|  | k151217_3 | strept | PV+,<br>vGAT+ | 0.6 | female | 19 | right | frontal | subcortical neoplasia |
|  | k151217_2 | strept | PV and<br>vGAT<br>non<br>concl | 0.65 | female | 19 | right | frontal | subcortical neoplasia |
|  | k121217_1 | strept | PV+,<br>vGAT+ | 0.5 | female | 68 | right | temporal | hydrocephalus |
|  | k080917_1 | NA |  | 0.5 | male | 45 | right | fronto-<br>temporal | subcortical neoplasia |
|  | k040915_2 | strept | PV non<br>concl | 0.5 | male | 55 | right | frontal | subcortical neoplasia |
|  | k050615_1 | strept | vGAT+ | 0.5 | male | 68 | right | temporal | glioblastoma grade IV. |
|  | k100417_1 | strept | PV+,<br>vGAT+ | 0.4 | female | 39 | right | frontal | glioblastoma grade IV. |
|  | k140415_2 | strept | vGAT<br>non<br>concl | 0.4 | male | 18 | right | frontal | malignant germinoma, grade II. |
|  | k190315_1 | strept | PV non<br>concl | 0.4 | male | 85 | left | fronto-<br>temporal | hydrocephalus |
|  | k230615_1 | strept | vGAT+ | 0.4 | female | 30 | no info | no info | hydrocephalus |
|  | k230915_1 | strept | PV non<br>concl | - | female | 40 | right | frontal | anaplastic ependymoma |
|  | k230915_3 | strept | vGAT+ | 0.45 | female | 40 | right | frontal | anaplastic ependymoma |
|  | k250615_1 | strept | PV+ | 0.45 | female | 10 | left | frontal | pilocytic astrocytoma |
|  | k250615_5 | strept | PV+ | 0.4 | female | 10 | left | frontal | pilocytic astrocytoma |
|  | k260515_1 | strept | NA | 0.4 | female | 31 | left | temporo-occipital | fibrillary astrocytoma |
|  | k280415_1 | strept | PV+,<br>vGAT+ | 0.4 | female | 28 | right | temporal | hydrocephalus |
|  | k280815_1 | strept | PV+ | 0.4 | female | 58 | right | temporal | subarachnoid hemorrhage |
|  | k290415_2 | non<br>concl |  | 0.4 | female | 32 | left | frontal | subcortical neoplasia |
| **pvBC-pvBC**<br>**pairs** |  |  |  |  |  |  |  |  |  |
|  | K151217_3<br>presynaptic | strept | PV+,<br>vGAT+ | 0.6 | female | 19 | right | frontal | subcortical neoplasia |
|  | postsynaptic | NA |  | 0.5 |  |  |  |  |  |
|  | k190619_1<br>mutual pair |  |  |  |  |  |  |  |  |
|  | cell 1 | NA |  | 0.55 | female | 56 | left | temporal | subcortical neoplasia |
|  | cell 2 | NA |  | 0.6 |  |  |  |  |  |
| **Fig 2 exps**<br>**mouse** |  |  |  |  |  |  |  |  |  |
|  | K250618_1 | strept |  | 0.4 |  |  |  |  |  |
|  | k260618_2 | strept |  | 0.5 |  |  |  |  |  |
|  | k260618_5 | NA |  | 0.5 |  |  |  |  |  |
|  | K020718_3 | strept |  | 0.4 |  |  |  |  |  |
|  | K030718_1 | strept |  | 0.55 |  |  |  |  |  |
|  | K030718_3 | NA |  | 0.7 |  |  |  |  |  |
|  | K290618_2 | strept | PV+ | 0.6 |  |  |  |  |  |
|  | K290618_3 | strept | PV+ | 0.5 |  |  |  |  |  |
|  | K290618_4 | strept |  | 0.5 |  |  |  |  |  |
|  | V260618_1 | NA |  |  |  |  |  |  |  |
|  | V290618_4 | strept | PV+ | 0.7 |  |  |  |  |  |
| **Fig 4 exps** |  |  |  |  |  |  |  |  |  |
|  | v101118_1 | strept |  | 0.6 | male | 20 | right | frontal | hydrocephalus |
|  | v15_02_18_2 | strept |  | 0.7 | female | 67 | right | occipital | subcortical neoplasia |
|  | v15_02_15_3 | strept |  | 0.7 | female | 67 | right | occipital | subcortical neoplasia |
|  | V26_02_19_2 | strept |  | 0.6 | male | 68 |  | temporal | hydrocephalus |
|  | V091118_1 | NA |  | 0.6 | female | 81 | right | temporal | hydrocephalus |
|  | V050419_2 | strept |  | 0.5 | female | 22 | right | frontal | hydrocephalus |
|  | v290319_2 | NA |  | 0.5 | male | 51 | no info | no info | hydrocephalus |
| NonFS | V260219_1 | strept |  | 1 | male | 68 |  | temporal | hydrocephalus |
|  | V120219_1 | strept |  | 0.8 | female | 66 | right | frontal | astrocytoma |
|  | V080119_2 | strept | vGAT+ | 0.9 | male | 49 | right | frontal | colloid cyst |
|  | v080119_1 | strept | vGAT+ | 0.8 | male | 49 | right | frontal | colloid cyst |
|  | V160119_1 | strept |  | 0.85 | male | 58 | left | frontal | epithelial carcinoma metaplasia |
|  | V131218_1 | strept |  | 0.9 | female | 79 | right | temporal | hydrocephalus |
|  | V050419_1 | strept |  | 0.85 | female | 22 | right | frontal | hydrocephalus |
|  | V13_12_18_1 | strept |  | 0.9 | female | 79 | right | temporal | hydrocephalus |
| **Other nonFS reported in results not in figures** |  |  |  |  |  |  |  |  |  |
|  | v190918_3 |  |  | 0.9 | male | 21 | right | temporo-occipital | subcortical cystic neoplasm |
|  | v311017_2 |  |  | 0.8 | female | 39 | right | frontal | glioblastoma |
|  | v091117_3 |  |  | 1.2 | female | 63 | right | temporal | hydrocephalus |
|  | v100818_1 |  |  | 0.9 | female | 46 | right | temporal | glioma (astrocytoma) |
|  | v130618_1 |  |  | 1.1 | female | 35 | right | temporal | subcortical neoplasia |
|  | v100518_2 |  |  | 1 | female | 73 | right | frontal | biopsy (MRI: multifocal lesions) |
|  | v220218_1 |  |  | 0.85 | male | 42 | left | frontal | subcortical neoplasia |
|  | v091117_2 |  |  | 0.9 | female | 63 | right | temporal | hydrocephalus |
|  | v231117_4 |  |  | 0.8 | female | 68 | no info | no info | hydrocephalus |
|  | v151117_1 |  |  | 0.9 | female | 61 | right | frontal | fibrillic astrocytoma |
|  | K300119_1 |  |  | 0.7 | female | 69 | no info | no info | hydrocephalus |
|  | K260219_2 |  |  | 0.7 | male | 68 |  | temporal | hydrocephalus |
|  | k231117_1 |  |  | 0.8 | female | 68 | no info | no info | hydrocephalus |
|  | k181017_1 |  |  | 0.8 | female | 78 | right | frontal | hydrocephalus |
| **Fig 5 exps** |  |  |  |  |  |  |  |  |  |
|  | v190619_2 | strept | PV+ | 0.6 | female | 56 |  |  |  |
|  | v190619_3 | strept | PV+ | 0.55 | female | 56 |  |  |  |
|  | v190619_4 | strept | PV+ | 0.6 | female | 56 |  |  |  |

**Supplementary table 2.** Recorded parameters used in computational simulation of human pvBCs.

| cell code | g leak (nS) | peak g autapse (nS) | peak g AHP (nS) | capacitance (pF) | AHP time to peak (ms) | AHP tau decay (ms) | AHP ascending tau (ms) | aut time of peak (ms) | aut decay tau (ms) |
| --- | --- | --- | --- | --- | --- | --- | --- | --- | --- |
| k260118_1 | 4.58 | 15.8 | 12.02 | 23.16 | 0.61 | 6.81 | 0.3 | 1.6 | 3.4 |
| k260118_2 | 2.69 | 9.61 | 7.40 | 23.92 | 1.07 | 4.94 | 0.3 | 1.8 | 4.0 |
| k080917_1 | 3.84 | 4.52 | 6.75 | 28.56 | 1.02 | 7.52 | 0.4 | 2.5 | 5.1 |
| k200218_6 | 8.16 | 6.13 | 13.80 | 80.75 | 0.52 | 6.21 | 0.3 | 3.0 | 4.3 |
| k121217_3 | 6.63 | 5.32 | 17.96 | 52.02 | 0.58 | 3.37 | 0.3 | 2.5 | 3.2 |
| k151217_3 | 2.62 | 7.38 | 28.94 | 24.93 | 0.88 | 2.90 | 0.4 | 2.5 | 5.2 |

